# Spatial Distribution of Breast Cancer in Sudan 2010-2016

**DOI:** 10.1101/516062

**Authors:** Marwa Maweya Abdelbagi Elbasheer, Ayah Galal Abdelrahman Alkhidir, Siham Mohammed Awad Mohammed, Areej Abuelgasim Hassan Abbas, Aisha Osman Mohamed, Isra Mahgoub Bereir, Hiba Reyad Abdalazeez, Mounkaila Noma

**Author notes:** Corresponding author (MMAE).

## Abstract

**Background:** Breast cancer is the most prevalent cancer among females worldwide including Sudan. The aim of this study was to determine the spatial distribution of breast cancer in Sudan.

**Materials and methods:** A facility based cross-sectional study was implemented in eighteen histopathology laboratories distributed in the three localities of Khartoum State on a sample of 4630 Breast Cancer cases diagnosed during the period 2010-2016. A master database was developed through Epi Info™ 7.1.5.2 for computerizing the data collected: the facility name, type (public or private), and its geo- location (latitude and longitude). Personal data on patients were extracted from their respective medical records (name, age, marital status, ethnic group, State, locality, administrative unit, permanent address and phone number, histopathology diagnosis). The data was summarized through SPSS to generate frequency tables for estimating prevalence and the geographical information system (ArcGIS 10.3) was used to generate the epidemiological distribution maps. ArcGIS 10.3 spatial analysis features were used to develop risk maps based on the kriging method.

**Results:** Breast cancer prevalence was 3.9 cases per 100,000 female populations. Of the 4423 cases of breast cancer, invasive breast carcinoma of no special type (NST) was the most frequent (79.5%, 3517/4423) histopathological diagnosis. The spatial analysis indicated as high risk areas for breast cancer in Sudan the States of Nile River, Northern, Red Sea, White Nile, Northern and Southern Kordofan.

**Conclusions:** The attempt to develop a predictive map of breast cancer in Sudan revealed three levels of risk areas (risk, intermediate and high risk areas); regardless the risk level, appropriate preventive and curative health interventions with full support from decision makers are urgently needed.

## Introduction

Breast cancer (BC) is a heterogeneous group of diseases characterized by different pathologies, biological characteristics and clinical behaviors. It is the leading cancer among females worldwide with 641,000 cases reported in 1980 and 1,643,000 cases in 2010; the annual incidence increased between the two years was 3.1% [1]. In the year 2012, globally, BC represented 25% of all cancers and 15% of the cancer deaths among females [2]; in 2015, WHO reported 571000 deaths [3] while by 2020, 1.7 million new cases are expected mostly in the developing countries [4]. The recent shift in its burden in the developing world is revealed by a high mortality rate and a poorer overall survival [2, 4]. The geographical distribution of BC in Africa revealed a marked variation in incidence within the continent with a high incidence rate of 130 cases/100,000 populations in Northern African countries and a lowest rate of 95 cases/100,000 populations recorded in the Western part of the African continent [5]. The highest standardized mortality rate worldwide according to WHO six regions was found in the East Mediterranean Office (EMRO) and Africa Regional Office (AFRO) with respectively 18.6% and 17.2% [6].

In Sudan, the burden of cancer had increased from 303 cases in 1967 to 6303 in 2010 in which the BC represented the most common cancer [7]. Further studies [8,9] reported that the highest prevalence of cancers was recorded in the States of Khartoum, North Kordufan, Nile River, Northern, Gezira and White Nile states and BC was the most prevalent. According to the records of the Radiation Isotope Center Khartoum (RICK) and Gezira Institute for Cancer treatment and Molecular Biology (GICMB), BC was the most predominant malignancy among females with respectively 29-34.5% and 30.0% of the cancers registered. Most cases were young aged women. About 40% were below 45 years (mean age of 50) with late advanced disease. On the other hand, male cancer constituted 3.5-4% [10, 11]. Furthermore, studies from Red Sea State (2003-2006) and Central Sudan (1999-2006) revealed that the majority of the patients were premenopausal women (age <50 years) who presented with a late stage metastasized disease [12, 13].

A study [14] was conducted based on 6771 cases of cancers diagnosed in Khartoum State by Sudan First National Cancer registry during the period 2009 to 2010. The findings revealed that the most common cancer was Breast cancer with an incidence rate of 25.1 per 100,000. The study also reported the possibility of underestimation of the burden which could be due to factors such as stigmatization and poverty, leading to undiagnosed or untreated cases. Overestimation was also pointed out for elderly patients who might be treated symptomatically at primary care levels or died before reaching cancer specialized institutions.

Available statistics on breast cancer in Sudan are mostly restricted to central institutions such as RICK and GICMB and the geographical distribution of the disease yet is unknown. This paper aimed to estimate the burden of breast cancer and provide its spatial distribution country-wide.

## Materials and methods

A facility based cross-sectional study was implemented. Data were extracted from eighteen histopathology laboratories within Khartoum State (Fig 1). In each of these laboratories, data collected included facility name, type (public or private), and geo-location (latitude and longitude). Personal data extracted from the facility records were name, age, marital status, ethnic group, state, locality, administrative unit, permanent address and phone number. Other information obtained from the records included the date of diagnosis and the histopathology diagnosis.

**Fig 1:**
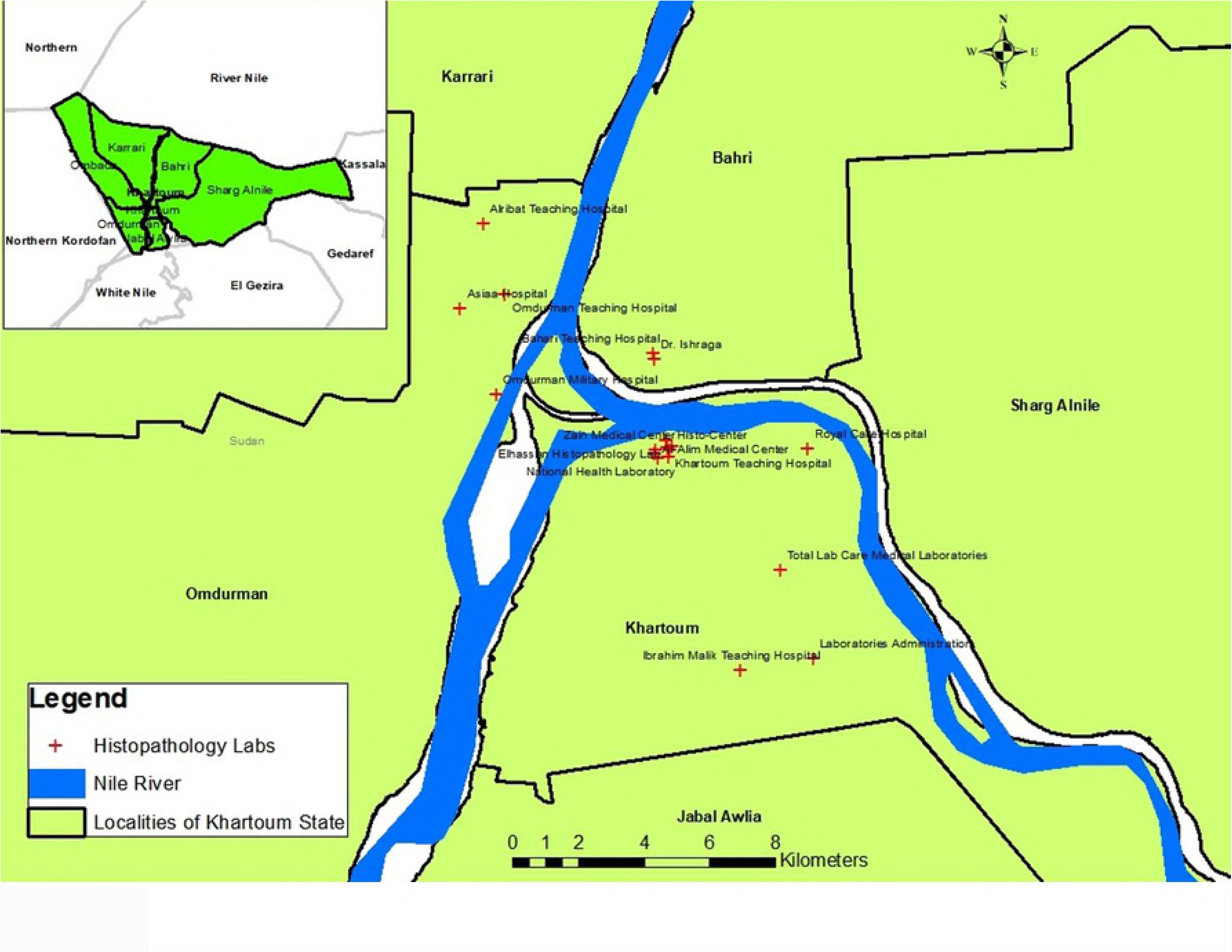
Geographical distribution of the histopathology laboratories

The master database, consisting of 4630 patient medical records was developed through Epi Info™ 7.1.5.2 and thereafter cleaned through the statistical package for social sciences (SPSS version 23) to exclude cases lacking important information such as the histopathology diagnosis, and date of diagnosis, as well as duplicated cases which were entered twice. The data of the remaining 4423 records was then summarized through SPSS to generate the frequency distribution of the cases in term of person (age, gender) and type of cancer diagnosed by the histopathology centers. Histopathology diagnoses recorded were invasive ductal carcinoma, invasive lobular carcinoma, carcinoma in situ and others, which were then regrouped to fit WHO 2012 classification [15]. The epidemiological distribution of breast cancer in Sudan was based on 1135 records for which data on residence were available. Those 1135 records were geo-referenced to facilitate the plotting of the residence of the patients. Prevalence was estimated using the updated 2016 Sudan Census Bureau and Statistics population data as a reference. ArcGIS 10.3 spatial analysis features were used to develop a risk map based on the krigging method [16].

## Ethical Statement

Study ethical clearance was obtained from Khartoum State Ministry of Health, Directorate of innovation and Scientific research Ethical Committee on 11th May 2017. (S1 file)

## Results

A total of 4423 cases of breast cancer were recorded (2010-2016) from eighteen laboratories distributed in Khartoum State. Patients were aged 12 to 103 years with an average (median) age of 48 years. They were predominately females 97.4% (4300/4413). The mean age at presentation was higher in males (61 years ±14.9) than in females (49 years ±14.2). Of the 4423 cases of breast cancer, invasive breast carcinoma of no special type (NST) was the most frequent histopathological diagnosis (79.5%, 3517/4423) followed by special subtypes of invasive carcinoma (12.4%, 547/4423) and precursor lesions (3.2%, 142/4423) and the remaining 4.9% were classified as others. Females were paying the highest burden with a crude prevalence of 3.9 cases per 100,000 female populations, ranging from 0.3 (Gedaref and Western Kordofan) to 22.1 in Khartoum as shown by Table 1 and Fig 2. On the other hand, male breast cancer was <1 per 100,000 male populations.

**Table 1:** Crude Prevalence (cases/100,000 population) of Breast Cancer in Sudan, data from eighteen histopathology laboratories located in Khartoum State (n=1135)

| States | Cases recoded from Labs |  |  | Prevalence per 100,000 population* |  | Total population |  | SCBS |
| --- | --- | --- | --- | --- | --- | --- | --- | --- |
|  | Number | Female | Male | Female | Male | Female | Male | 2016 |
| Khartoum | 786 | 766 | 20 | 22.1 | 0.5 | 3,464,536 | 3,920,622 | 7,385,158 |
| Red Sea | 79 | 77 | 2 | 12.5 | 0.2 | 616,423 | 828,930 | 1,445,353 |
| El Gezira | 45 | 44 | 1 | 1.8 | 0.1 | 2,464,166 | 2,295,598 | 4,759,764 |
| Northern Kordofan | 45 | 44 | 1 | 2.7 | 0.1 | 1,628,070 | 1,512,107 | 3,140,177 |
| White Nile | 43 | 42 | 1 | 3.5 | 0.1 | 1,183,915 | 1,140,529 | 2,324,444 |
| River Nile | 30 | 29 | 1 | 4.2 | 0.1 | 699,980 | 729,533 | 1,429,513 |
| Southern Darfur | 27 | 26 | 1 | 2.1 | 0.1 | 1,244,275 | 1,299,942 | 2,544,217 |
| Northern | 20 | 19 | 1 | 4.4 | 0.1 | 438,160 | 448,851 | 887,011 |
| Sennar | 17 | 17 | 0 | 1.8 | 0.1 | 912,230 | 865,752 | 1,777,982 |
| Northern Darfur | 14 | 14 | 0 | 1.2 | 0.0 | 1,115,490 | 1,165,395 | 2,280,885 |
| Blue Nile | 7 | 7 | 0 | 1.3 | 0.0 | 517,492 | 531,874 | 1,049,366 |
| Southern Kordofan | 6 | 6 | 0 | 1.2 | 0.0 | 501,841 | 490,106 | 991,948 |
| Western Darfur | 6 | 6 | 0 | 2.0 | 0.1 | 285,234 | 273,108 | 558,342 |
| Kassala | 5 | 5 | 0 | 0.5 | 0.0 | 1,053,571 | 1,306,512 | 2,360,083 |
| Gedaref | 3 | 3 | 0 | 0.3 | 0.0 | 1,012,329 | 999,285 | 2,011,614 |
| Western Kordofan | 1 | 1 | 0 | 0.3 | 0.0 | 282,473 | 275,868 | 558,342 |
| Rumbeck** | 1 |  |  |  |  |  |  |  |
| **Total** | **1135** | **1105** | **29** | **3.9** | **0.1** | **17,420,186** | **18,084,011** | **35,504,197** |

SCBS: Sudan Census Bureau and Statistics
\* Crude prevalence computed as number of breast cancer cases/ total female population x 100,000 female population all age
\*\* In Lakes State of South Sudan

**Fig 2:**
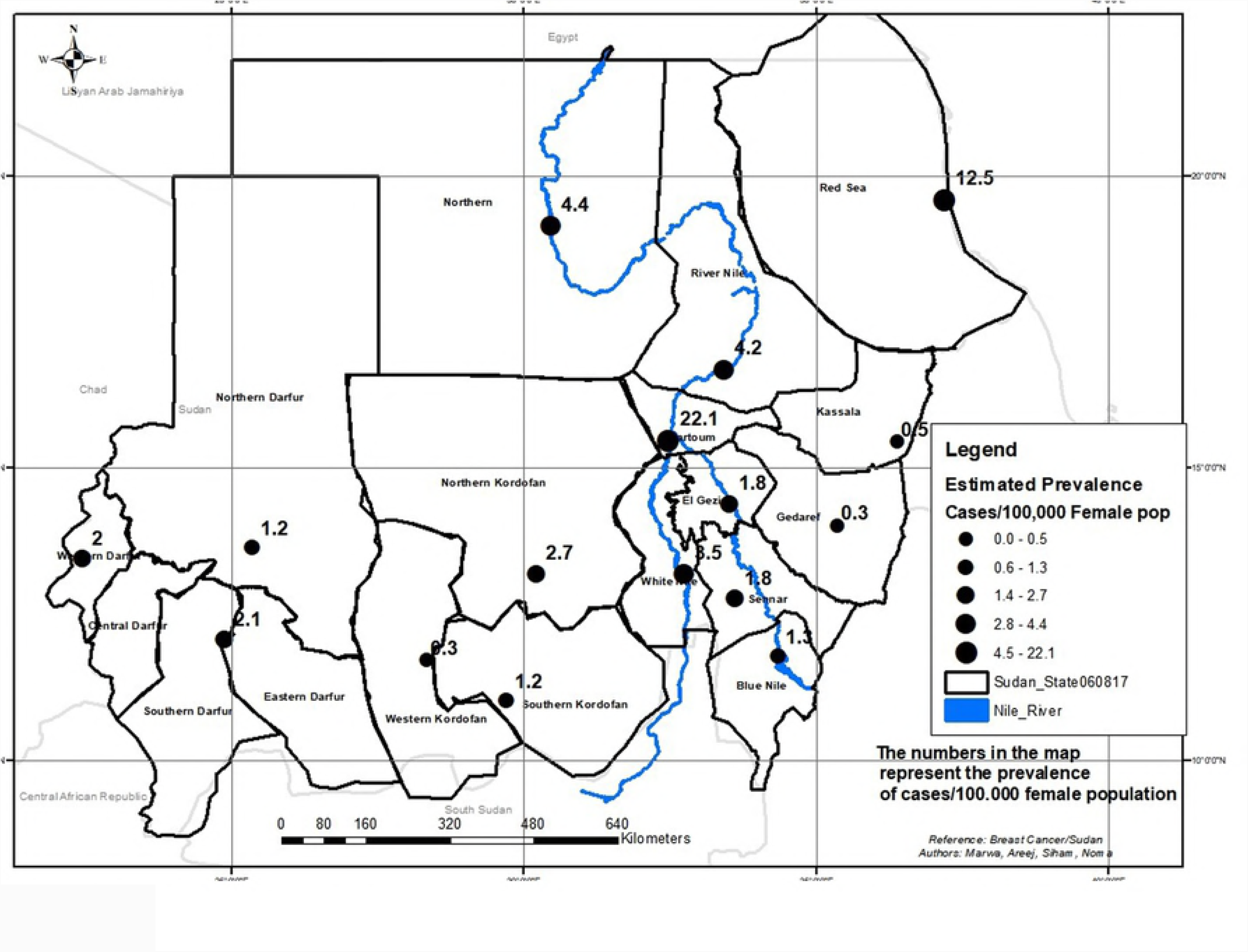
Epidemiological distribution of Breast Cancer in female population in Sudan (n=1135)

The spatial analysis confirmed that Breast Cancer is a country-wide health problem. The risk of breast cancer according to the map generated using the kriging method of the spatial distribution indicated three gradient scale colors of risk (Fig 3). Risk areas included Western, Central, Southern Darfur and partially Northern states, and a large part of Red Sea; Invasive carcinoma was predominant type in those States. Intermediate risk areas, a mosaic for invasive carcinoma NST, Special Subtypes of Invasive Carcinoma and Precursor lesions, included the States of Khartoum, Gezira, White Nile, Kassala, Gedaref, Sennar, Eastern Darfur, and focally Northern Darfur. High risk areas were the States of Nile River, Northern, Red Sea (focal), White Nile, Northern and Southern Kordofan.

**Fig 3:**
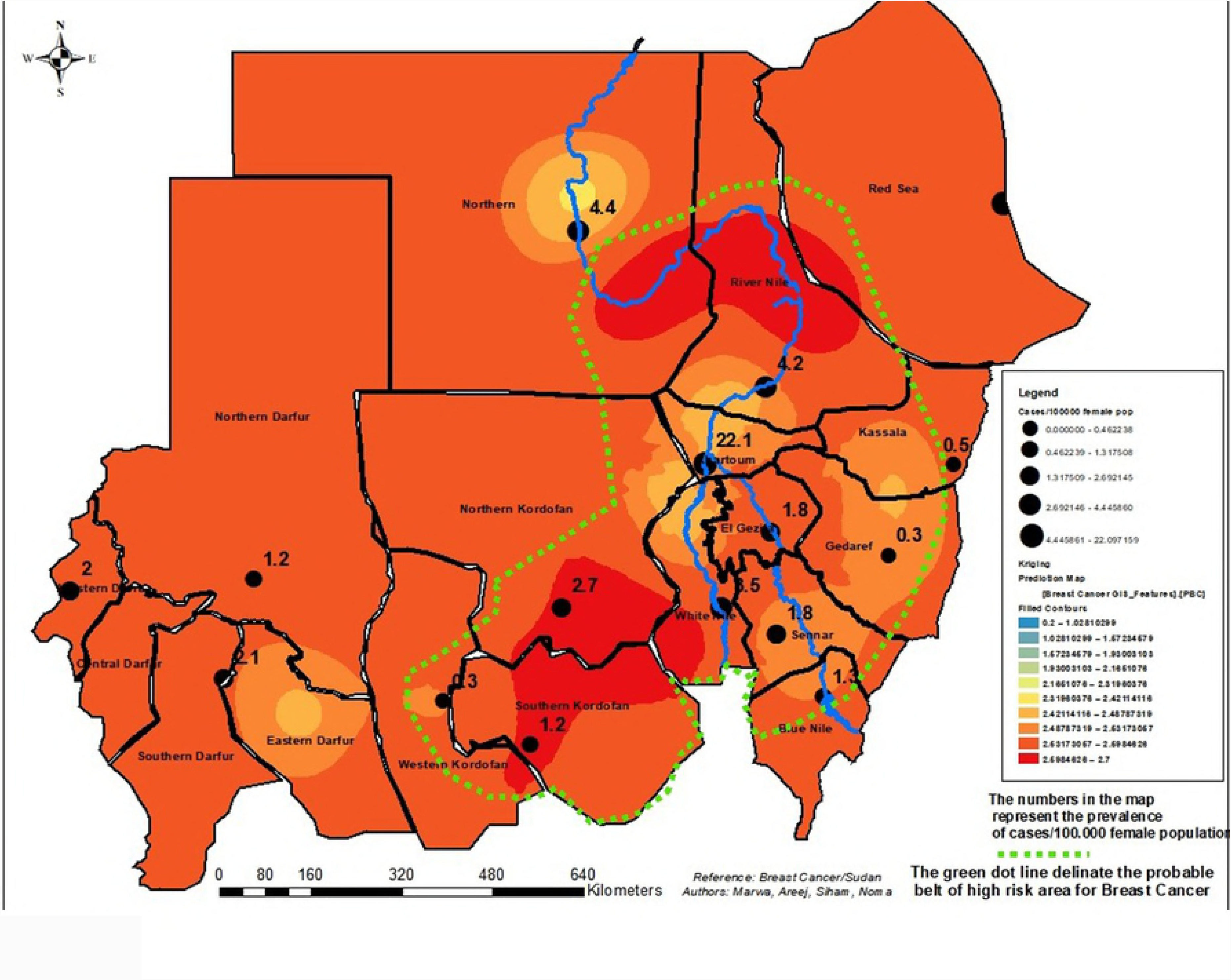
Breast Cancer risk map in Sudan (n=1135)

## Discussion

Our findings revealed a crude prevalence of 3.9 cases per 100,000 female populations for the period 2010 to 2016. This burden of Sudan females was also reported elsewhere in Sub-Saharan African countries, fluctuating from 4.5% (Zimbabwe) to 38.9/100 000 females in (South Africa) [17].

The average age of our patients was 48 years revealing that younger population was affected as reported in Central Africa (45.83 years) and Ghana (49.1 years) [18, 19]. On the contrary in developed countries, women are affected at older age respectively at 57 years and 62 years in New Zealand and United States [20, 21].

Invasive carcinoma of NST was the prevalent type (79.5%) of breast cancer in our study as previously published in Sudan [10], elsewhere it was 60% of breast cancer cases as reported by Badowska-Kozakiewicz, et al. [22].

The Epidemiological map generated per states indicated that the highest prevalence was recorded in Khartoum and Red Sea States with respectively 22.1 and 12.5 per 100,000 female populations. The figures from Khartoum and Red Sea States may be interpreted as related to the fact that most of the cases were reported from those States. We endorse the contrary based on the modeling which revealed that the spatial distribution predicts Khartoum State as intermediate risk whereas Red Sea State was displayed with a highly focal risk area. In the overall, we would like to emphasis that the breast cancer is a country wide public health problem. The delineated belt is a subject for further discussion related to the modeling technique and the limitations of the data which not include environmental and socio-economic factors.

This rapid evidence based delineation of breast cancer areas is a tool for guiding public health professionals and decision makers to establish a breast cancer program for fine tuning the epidemiological map and the subsequent risk map generated as applied in health sector in Iran [23] and elsewhere in Saudi Arabia [24] where the geographical information was used to set priorities. One of the limitations of our model is the lack of environmental data to better assess the pattern of breast cancer which is a multi-factorial condition. The risk map was developed based on individual location of residence reported by the patients which may be a limiting factor leading to the over estimation or under estimation of the number of cases per State. This potential bias according to us was triggered out by the modeling approach of the krigging method which was used.

## Conclusion

Our findings provided an understanding of the pattern of the spatial distribution of breast cancer country wide with hot spots defined as high risk and intermediate risk areas. As further data may be needed to improve the risk map, the decision makers and the health professionals should for equity reasons look at decentralizing of the health system which could not be efficient and operational if all the expertise are concentrated mainly in the State hosting the capital of country.

## Acknowledgements

Authors would like to acknowledge the administration and staff of the eighteen histopathology laboratories for permitting the extraction of the data.

## Supporting Information

**S1 File: Ethical Clearance Certificate**

